# Differential Effects of Cocaine and Morphine on the Diurnal Regulation of the Mouse Nucleus Accumbens Proteome

**DOI:** 10.1101/2023.03.01.530696

**Authors:** Kyle D. Ketchesin, Darius D. Becker-Krail, Xiangning Xue, Rashaun S. Wilson, TuKiet T. Lam, Kenneth R. Williams, Angus C. Nairn, George C. Tseng, Ryan W. Logan

## Abstract

Substance use disorders (SUDs) are associated with disruptions in sleep and circadian rhythms that persist during abstinence and may contribute to relapse risk. Repeated use of substances such as psychostimulants and opioids may lead to significant alterations in molecular rhythms in the nucleus accumbens (NAc), a brain region central to reward and motivation. Previous studies have identified rhythm alterations in the transcriptome of the NAc and other brain regions following the administration of psychostimulants or opioids. However, little is known about the impact of substance use on the diurnal rhythms of the proteome in the NAc. We used liquid chromatography coupled to tandem mass spectrometry-based (LC-MS/MS) quantitative proteomics, along with a data-independent acquisition (DIA) analysis pipeline, to investigate the effects of cocaine or morphine administration on diurnal rhythms of proteome in the mouse NAc. Overall, our data reveals cocaine and morphine differentially alters diurnal rhythms of the proteome in the NAc, with largely independent differentially expressed proteins dependent on time-of-day. Pathways enriched from cocaine altered protein rhythms were primarily associated with glucocorticoid signaling and metabolism, whereas morphine was associated with neuroinflammation. Collectively, these findings are the first to characterize the diurnal regulation of the NAc proteome and demonstrate a novel relationship between phase-dependent regulation of protein expression and the differential effects of cocaine and morphine on the NAc proteome.

## INTRODUCTION

The ongoing opioid use disorder (OUD) and substance use disorder (SUD) epidemic continues to worsen in the United States and globally, exacerbated by the recent COVID19 pandemic. In 2022, the United States witnessed the highest number of deaths from drug overdose ever recorded ^1^, attributed to increased use of opioids and other substances. While mainstay treatments such as methadone and buprenorphine are effective, attention remains on identifying new targets for therapeutic development through research investigating the consequences of opioids on molecular pathways in the brain. A key brain region involved in the regulation of reward, motivation, and mood, associated with OUD, is the nucleus accumbens (NAc) ^2^. A majority of the work investigating the impact of substance use on the NAc focuses on either specific genes or proteins of interest, or transcriptomic changes ^3–5^. However, studies investigating the impact of substance use on the proteomic landscape are necessary in the brain, as alterations in protein expression may be highly reflective of drug-induced changes in molecular and cellular functions ^6–8^. Notably, few studies have used quantitative proteomics to investigate the effects of psychostimulants and opioids in brain ^9–15^, while no studies, to date, have involved the diurnal rhythm regulation of the proteome in NAc. Recent work from us and others find molecular rhythms in NAc and other brain regions regulate substance reward, craving, and relapse, highlighting critical roles for sleep and circadian rhythms in SUDs ^16–26^.

In mammals, circadian rhythms are generated and maintained by the suprachiasmatic nucleus (SCN), an endogenous autonomous timekeeping nucleus within the anterior hypothalamus of the brain ^27^. The SCN serves as the core pacemaker, relaying light-entrained temporal information to synchronize rhythms in physiology at the system, tissue, cellular, and molecular levels ^27,28^. While the SCN is considered the central pacemaker, extra-SCN oscillators are present throughout the brain, including NAc and other reward-related regions ^29,30^, resembling diurnal rhythmicity of neuronal activity, and gene and protein expression and function. Molecular rhythms are regulated by a series of transcriptional and translational feedback loops, oscillating on a near 24-hour timescale (reviewed in Partch *et al.* ^31^). Disruptions to rhythms in neuronal activity and/or molecular rhythms in NAc are associated with altered mood, anxiety, and reward-related behaviors ^18,20,21,32–39^. Importantly, psychostimulants and opioids are shown to alter transcriptional rhythms in the NAc of both humans and rodents ^18,19,32,40–48^, further suggesting drug-induced changes in diurnal regulation of molecular rhythms functionally impacts brain physiology and behavior related to substance use. Identifying the effects of substance use on the NAc proteome is a necessary first step to advance our understanding of the role of circadian rhythms in the etiology and treatment of SUDs.

In the present study, we used liquid chromatography coupled to tandem mass spectrometry-based (LC-MS/MS) quantitative proteomics to investigate the effects of the psychostimulant, cocaine, or the opioid, morphine, on regulation of the proteome in NAc of mice. We employed a data-independent acquisition (DIA) analysis pipeline ^49,50–53^, affording greater reproducibility, sensitivity, and dynamic range compared to traditional data dependent acquisition (DDA) approaches ^49,50–53^. Notably, our approach assessed diurnal rhythms of the proteome in mouse NAc across multiple timepoints following cocaine or morphine exposure, capturing both differentially expressed proteins and the impact of substance use on the temporal variation of the proteome ^51,53^.

## MATERIALS AND METHODS

### Animals

Male C57BL/6J mice (The Jackson Laboratory; Bar Harbor, ME; IMSR Cat# JAX:000664, RRID:IMSR_JAX:000664), ages 8-12 weeks, were maintained on a 12:12 light-dark cycle (lights on at 0700, zeitgeber time (ZT) 0, and lights off at 1900, ZT12). Food and water were provided *ad libitum*. Animal use and procedures were conducted in accordance with the National Institute of Health guidelines and approved by the University of Pittsburgh Institutional Animal Care and Use Committee.

### Drug Administration and Tissue Collection

Cocaine hydrochloride and morphine were provided by the National Institute on Drug Abuse (NIDA) and dissolved with 0.9% saline (Fisher Scientific). Mice were injected intraperitoneally (i.p.) for 7 days with either saline (10 ml/kg), cocaine (15 mg/kg at 10 ml/kg), or morphine (10 mg/kg at 10 ml/kg). Drugs were administered between ZT4-8, as previously described ^40^. Following the final injection (24-hours), 6 mice per treatment group were sacrificed via cervical dislocation at either ZT5, ZT10, ZT17, or ZT22, and brains were removed and rapidly frozen on dry ice. Brains were sectioned to collect bilateral punches from the NAc of each mouse, then immediately returned to −80C until further processing.

### Liquid Chromatography Mass Spectrometry (LC-MS/MS) Processing and Analysis

#### Sample Preparation

Preparation of mouse NAc samples for LC-MS/MS was performed as follows: 50 µL of RIPA containing protease inhibitor (cat#87786, ThermoFisher Scientific) and phosphatase inhibitor (cat#78420, ThermoFisher Scientific) were added to each sample. Samples were sonicated (15s with two 0.5s pulses), then centrifuged (4°C at 14.6K rpm) for 10min. Supernatant was transferred to a new vial and protein were precipitated using MeOH:Chloroform:Water (4:1:3). Extracted proteins were transferred to new tube and volume was brought to 100µL with 8M Urea, 400mM ammonium bicarbonate. Protein was reduced with 8µL of 45mM DTT and incubated at 37°C for 30min. Protein was alkylated with 8µL of 100mM iodoacetamide (IAM), then incubated in the dark at room temperature for 30min. After diluting with water to bring the urea concentration to 2M, sequencing-grade trypsin (Promega, Madison, WI, USA) was added at a weight ratio of 1:20 (trypsin:protein) and incubated at 37°C for 16-hrs. Samples were desalted using C18 spin columns (The Nest Group, Inc., Southborough, MA, USA) and dried in a rotary evaporator. Samples were resuspended in 0.2% trifluoroacetic acid (TFA) and 2% acetonitrile (ACN) in water prior to LC-MS/MS analysis.

#### Data-independent acquisition (DIA)

DIA LC-MS/MS was performed using a nanoACQUITY UPLC system (Waters Corporation, Milford, MA, USA) connected to an Orbitrap Fusion Tribrid (ThermoFisher Scientific, San Jose, CA, USA) mass spectrometer. After injection, samples were loaded into a trapping column (nanoACQUITY UPLC Symmetry C18 Trap column, 180µm × 20mm) at a flow rate of 5 µL/min and separated with a C18 column (nanoACQUITY column Peptide BEH C18, 75µm × 250mm). The compositions of mobile phases A and B were 0.1% formic acid in water and 0.1% formic acid in ACN. Peptides were eluted with a gradient extending from 6% to 35% mobile phase B in 90min and then to 85% mobile phase B in another 15min at a flow rate of 300 nL/min and a column temperature of 37°C. The data were acquired with the mass spectrometer operating in a data-independent mode with an isolation window width of 25m/z. The full scan was performed in the range of 400–1,000m/z with “Use Quadrupole Isolation” enabled at an Orbitrap resolution of 120,000 at 200m/z and automatic gain control (AGC) target value of 4 × 10^5^. Fragment ions from each peptide MS2 were generated in the C-trap with higher-energy collision dissociation (HCD) at a collision energy of 28% and detected in the Orbitrap at a resolution of 60,000.

#### DIA Data analysis

DIA spectra were searched against a peptide library generated from DDA spectra using Scaffold DIA software (v.1.1.1, Proteome Software, Portland, OR, USA). Within Scaffold DIA, raw files were first converted to mzML format using ProteoWizard (v.3.0.11748). The samples were then aligned by retention time and individually searched against a *Mus musculus* proteome database exported from UniProt with a peptide mass tolerance of 10ppm and a fragment mass tolerance of 10ppm. The data acquisition type was set to “Non-Overlapping DIA”, and the maximum missed cleavages was set to 1. Fixed modifications included carbamidomethylation of cysteine residues (+57.02). Peptides with charge states between 2-3 and 6-30 amino acids in length were considered for quantitation, and the resulting peptides were filtered by Percolator (v.3.01) at a threshold False Discovery Rate (FDR) of 0.01. Peptide quantification was performed by EncyclopeDIA v. 0.6.12, and 6 of the highest quality fragment ions were selected for quantitation ^53^. Proteins containing redundant peptides were grouped to satisfy the principles of parsimony, and proteins were filtered at a threshold of 2 peptides per protein and an FDR of 1%. Data were median normalized between the samples to remove unwanted experimental variation. Out of 3047 proteins, 58 have missingness in more than 40% of samples and are removed. The missingness in the remaining proteins was imputed with K-nearest-neighbors using the impute.knn() function in R package impute. All analyses were performed based on log2 transformed values.

### Differential Expression Analysis

We analyzed the treatment effect of cocaine or morphine separately using the saline group as the reference. For each treatment group, we performed the differential expression (DE) analysis during daytime and nighttime separately. Specifically, daytime refers to ZT5 and ZT10, while nighttime refers to ZT17 and ZT22. The DE analyses were performed using the R package limma. Proteins were considered differentially expressed if p<0.05 and log_2_ fold change <-0.1375 or >0.1375 (fold change ±1.1 or 10% expression change).

### Rhythmicity Analysis by Treatment Group

For each treatment group, we performed rhythmicity analysis for each protein by regressing the expression level to a sinusoidal function of time: E(y)=Asin(f(t+p))+M, where y is the expression level, A is the amplitude, f=2π/24 is the frequency of circadian rhythmicity corresponding to a 24-hour period, t is the time when the sample was collected (5, 10, 17, 22 corresponding to ZT5, ZT10, ZT17, and ZT22), p is the phase and M is the rhythm-adjusted mean. Nonlinear optimization using Levenberg-Marquardt algorithm was used for the estimation of the parameters Â, 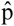, and 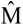. The goodness-of-fit coefficient R^2^ was used to assess the significance of rhythmicity. We calculate R^2^ using R^2^=1-RSS_m/RSS_0, where RSS_m is the residual sum of square of the fitted model and RSS_0 is the residual sum of square of the null model E(y)=M, thus R^2^ represents the percentage of total data variance explained by the rhythmicity model. The significance level of R^2^ was evaluated by permutation test. By shuffling the data, we disrupt the association between time and expression level and get a null distribution of R^2^. We shuffled the data 1000 times and pooled the null R^2^ from all proteins. The p-values of rhythmicity were calculated as the upper percentile of the observed R^2^ in the null R^2^ distribution, then we calculated the q-values using Benjamini-Hochberg procedure.

### Change of Rhythmicity Analysis between Treatment Group

The change of rhythmicity coefficient (including A, p, M and R^2^) between treatment groups was evaluated by permuting data between the two groups. Similar as the permutation for rhythmicity analysis, we generate a null distribution of ΔA, Δp, ΔM and ΔR^2^, and the p-values were calculated by comparing the observed differential parameter to its corresponding null distribution. Taking comparison between the morphine group and the saline group (using saline group as the baseline) as an example, proteins with ΔA > 0 has larger amplitude in morphine group compared to saline group; proteins with |Δp|>0 will peak at different time during a circadian cycle. These two parameters are only meaningful if the protein is rhythmic in both treatment groups. Proteins with |ΔM|>0 have a rhythm adjusted differential expression (DE). And proteins with ΔR^2^>0 have higher rhythmicity fitness in the morphine group. We restricted the comparison of R^2^ in proteins that are rhythmic in at least one group (i.e. a protein with gain of rhythmicity must be rhythmic in at least the morphine group and with the p-value of ΔR^2^ smaller than 0.05; a protein with loss of rhythmicity must be rhythmic in at least the saline group and with the p-value of ΔR^2^ smaller than 0.05).

### Enriched Pathways and Biological Processes Analysis

Both Ingenuity Pathway Analysis (IPA) software (QIAGEN; Hilden, Germany; RRID:SCR_008653)^54^ and the online bioinformatics database Metascape (https://metascape.org/; RRID:SCR_016620)^55^ were used to identify enriched molecular pathways and processes in our protein lists. For identification of significant pathways, a significance threshold of p<0.05 (or a −log10(p-value) of 1.3) was used. IPA software was used to identify enriched canonical molecular pathways (user dataset as the reference set), while Metascape was used to identify enriched biological processes using only Gene Ontology (GO) Biological Processes as the ontology source. Within Metascape, all statistically enriched terms, accumulative hypergeometric p-values, and enrichment factors were automatically calculated and used for filtering. The remaining significant terms were then hierarchically clustered into a tree based on Kappa-statistical similarities among their gene memberships, with a 0.3 kappa score applied as the threshold to cast the tree into term clusters. The most significant term in each cluster served as the cluster title. ^55^

## RESULTS

### Cocaine or morphine administration induces differential changes in the nucleus accumbens proteome across time of day

We used quantitative proteomics to investigate the effects of cocaine or morphine on the regulation of the mouse NAc proteome across time of day (Fig. 1A). DE proteins were detected between drug exposure and saline at both day (ZT5, ZT10) and night (ZT17, ZT22) (Fig. 1B; Datasets S1 & S2). Most of the DE proteins were detected in morphine exposed mice during the night (122 DE proteins). Notably, there was little overlap in DE proteins between substances and phase (Fig. 1C).

**Figure 1.**
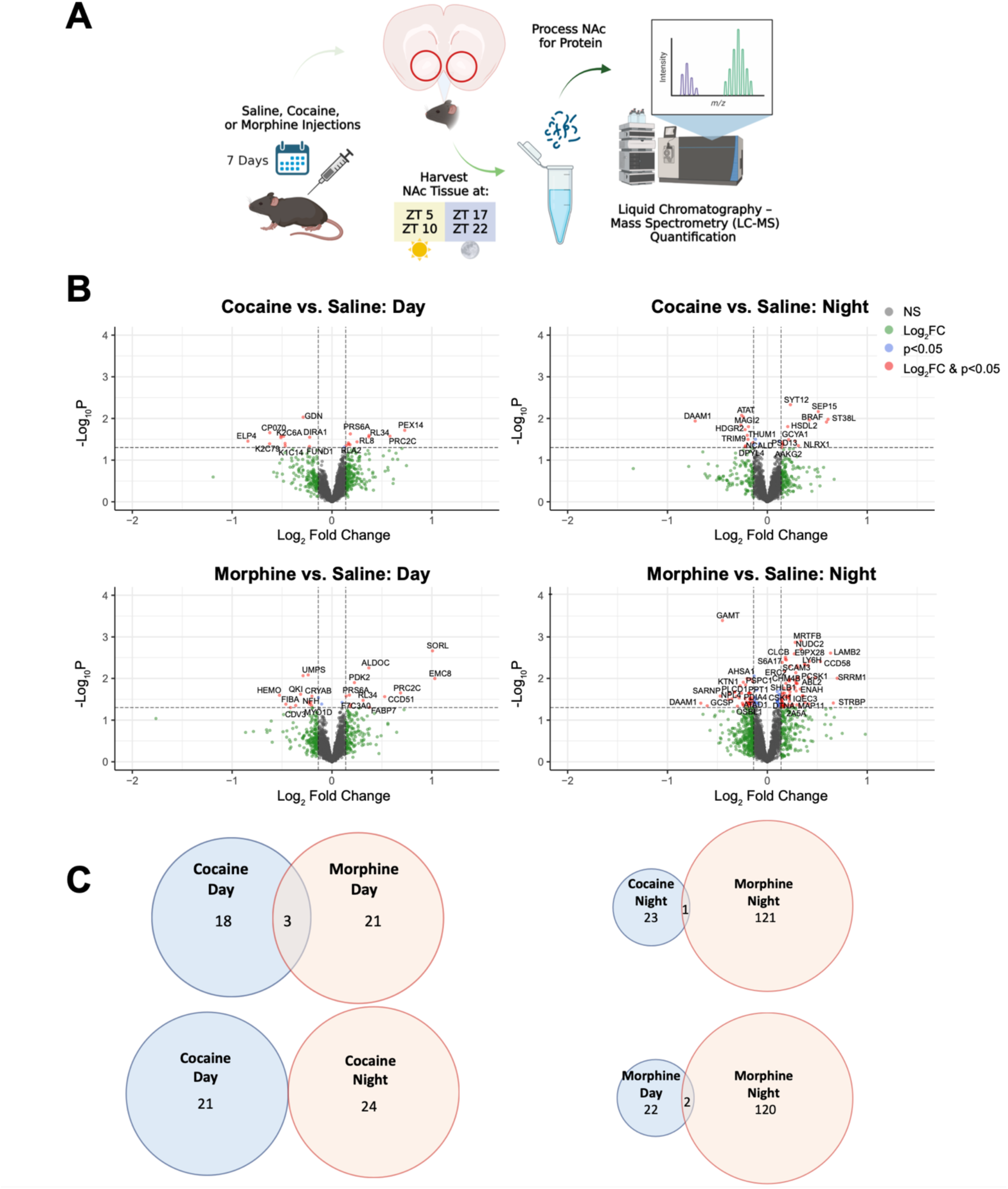
Cocaine or morphine induces differential changes in the nucleus accumbens proteome across time of day. (**A**) Schematic overview of the treatment paradigm and tissue collection prior to liquid chromatography mass spectrometry (LC-MS) quantification. Mice were injected intraperitoneally (i.p.) for 7 days with either saline (10 ml/k*g*), cocaine (15 mg/kg at 10 ml/kg), or morphine (10 mg/kg at 10 ml/kg). Following exposure, NAc tissue was collected across 4 times of day (ZT 5, 10, 17, & 22; ZT0 = 7am) and processed for LC-MS. (**B**) Volcano plots depicting differentially expressed (DE) proteins during the day (left; ZT 5 & 10) and night (right; ZT 17 & 22) as a result of cocaine (top) or morphine (bottom). Horizontal dashed line represents the p value significance cutoff (-Log_10_P of 1.3, or p=0.05), while the vertical dashed lines represent the fold change (FC) cutoff (Log_2_FC < −0.1375 or > 0.1375, or FC ± 1.1). Red dots indicate DE proteins that meet both cutoffs. (**C**) Venn diagrams depicting overlap of cocaine and morphine DE proteins during the day and night that met both FC and p value cutoffs. Venn diagram circles are sized by the number of proteins, proportional within comparisons. Schematic created with BioRender.com.

### Stress, immune, and metabolic-related pathways are enriched among the DE proteins in the NAc following cocaine or morphine administration

Next, we determined pathway and biological enrichment for DE proteins in the NAc following cocaine or morphine. DE proteins following cocaine administration were enriched for stress-related pathways (Figs. 2A & B; Datasets S3 & S4). For example, the top enriched pathways for the cocaine group were glucocorticoid receptor signaling during the day and corticotropin releasing hormone signaling during the night (Fig. 2A). Axonal-related pathways (e.g., semaphoring neuronal repulsive signaling) and processes (e.g., positive regulation of axonogenesis) were also enriched in cocaine groups during the night (Fig. 2B). Several of the top DE proteins related to glucocorticoid signaling were keratin proteins (e.g., KRT14, KRT5, KRT6B, KRT76, AND KRT79), which have been previously related to trauma and stress-related disorders ^56^. Following morphine administration, DE proteins were enriched for immune- and metabolic-related pathways (Figs. 2A & B; Datasets S3 & S4). For instance, metabolic-related pathways, such as LXR/RXR and FXR/RXR activation, were enriched during the day, and purine-containing compound metabolic processes were enriched for both day and night. Acute phase response signaling and CLTA4 signaling in cytotoxic T lymphocytes were enriched for day and night, respectively, following morphine.

**Figure 2.**
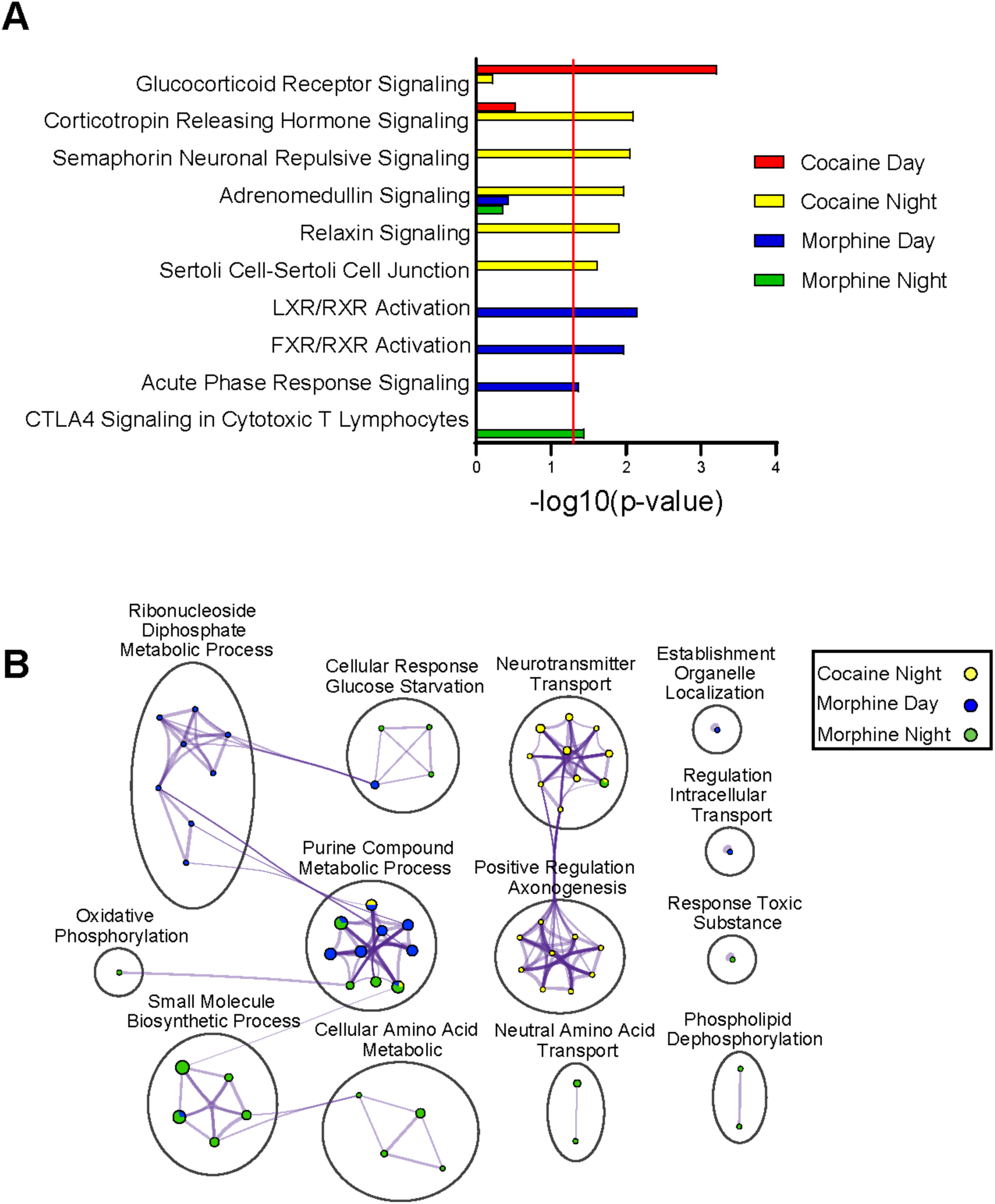
Pathway and biological process enrichment analysis for differentially expressed proteins in the NAc following cocaine or morphine. (**A**) Top pathways enriched among the DE proteins in the NAc by substance across time of day, as revealed by Ingenuity Pathway Analysis (IPA). A significance cutoff of −Log10(p value) of 1.3, or p<0.05, was used to determine pathway enrichment, depicted by the red line. (**B**) Gene Ontology (GO) Biological process enrichment via Metascape for the top DE proteins in the NAc by substance across time of day. Meta-analysis was used to compare process enrichment across substances by time of day, depicted via a Cytoscape enrichment network plot. Terms with p < 0.01, a minimum count of 3, and enrichment factor >1.5 were grouped into clusters based on their membership similarities. The cluster was named after the most statistically significant term within the cluster.). The nodes are represented as pie charts, where the size of the pie is proportional to the total number of proteins for that specific term, and color indicates the identity of the gene list, where the size of the slice represents the percentage of proteins enriched for each corresponding term. Similar terms are connected by purple lines (kappa score >0.3), and line thickness indicates degree of connectivity.

### Cocaine or morphine exposure alters diurnal rhythmicity in NAc proteome

We aimed to determine whether changes in rhythmicity contribute to differential protein expression in NAc. Using threshold-free approach to investigate overlap of proteins, RRHO analysis showed minimal overlap between cocaine- and morphine-specific rhythmic proteins in NAc (Fig. 3A; Dataset S5), suggesting psychostimulants and opioids lead to largely distinct alterations in rhythmic protein expression. We identified both gain and loss of rhythmicity in proteins following substance administration compared to saline treated mice (Fig. 3B). There was a striking gain of rhythmic proteins particularly following morphine, suggesting opioid-induced circadian reprogramming in NAc (Fig. 3B; representative scatterplots in Fig. 3C; Datasets 6 & 7). Like RRHO analysis (Fig. 3A), there is very little overlap between proteins that gain rhythmicity in morphine or cocaine (Fig. 3D; F6RPJ9 protein). In contrast, there was substantial overlap between proteins that lose rhythmicity following cocaine or morphine, suggesting similarities in the impact of these substances on the diurnal proteome in NAc.

**Figure 3.**
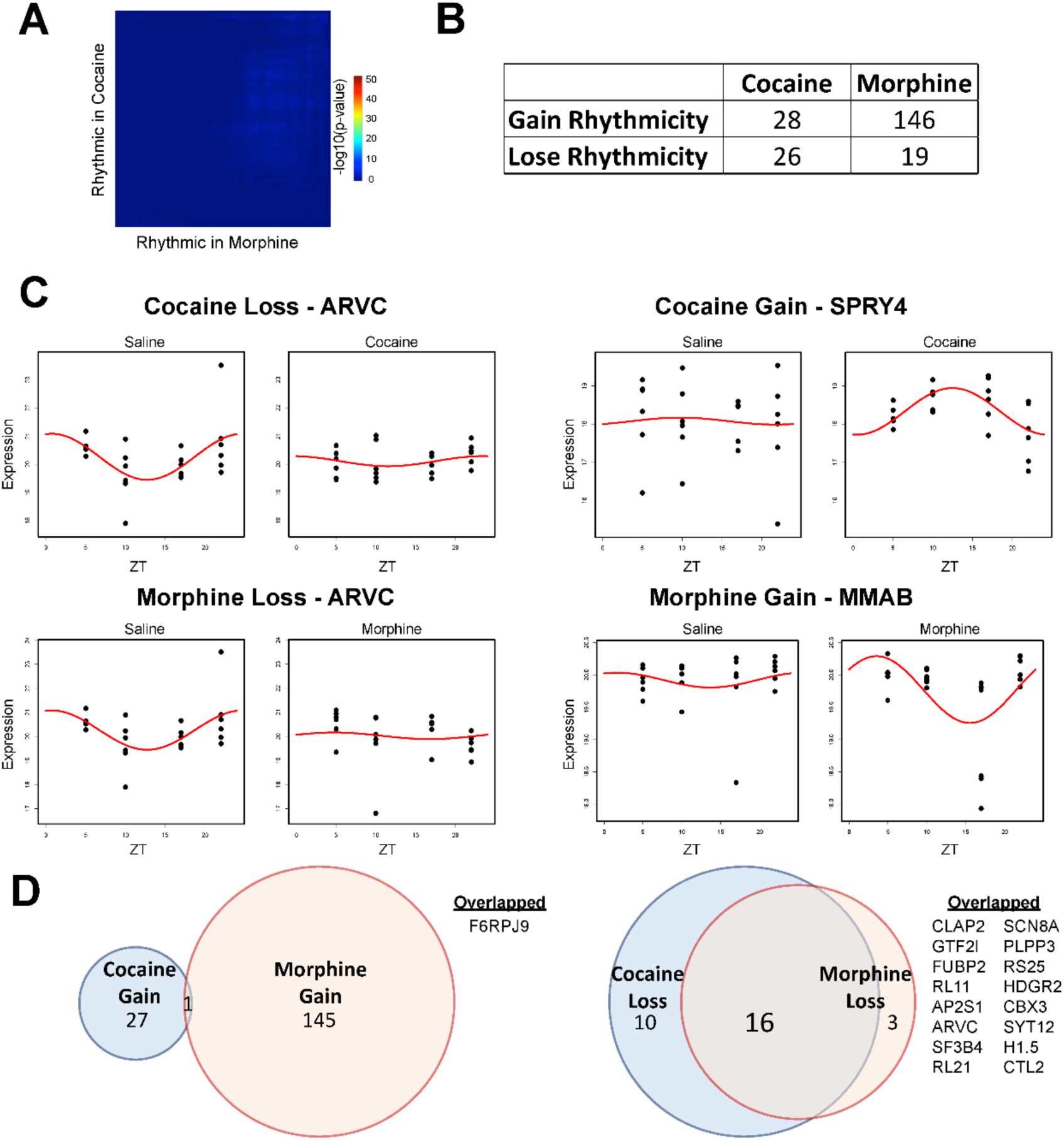
Cocaine or morphine alters rhythmicity in the NAc proteome. (**A**) Rank-Rank Hypergeometric Overlap (RRHO) analysis of the overlap between the cocaine and morphine specific rhythmic proteins in the NAc reveals limited overlap. Cosiner rhythmicity analysis was utilized to detect rhythmicity. (**B**) Table overview of the number of rhythmic proteins in the NAc that either gained or lost rhythmicity in the NAc, as a result of cocaine or morphine relative to saline treated mice. (**C**) Representative proteins with a gain (right) or loss of rhythmicity (left) in the NAc following cocaine (top) or morphine (bottom), relative to saline. Proteins with the greatest amplitude change were chosen. Dots indicate individual samples, with the y axis indicating expression and x axis indicating time of day (ZT 5, 10, 17, 22). Red line depicts the fitted curve as revealed in the cosiner rhythmicity analysis. (**D**) Venn diagrams depicting overlap of proteins that either gained (left) or lost (right) rhythmicity following cocaine or morphine. Overlapped proteins with gained or lost rhythmicity are listed on the right. Venn diagram circles are sized by the number of proteins, proportional within comparisons.

### Enrichment of protein translation, metabolic, and inflammation-related pathways in gain or loss of rhythmicity in NAc following cocaine or morphine

We determined enrichment of pathways for proteins that displayed a gain or loss of rhythmicity in NAc following cocaine or morphine. We found proteins related to translation (i.e., EIF2 signaling) lose rhythmicity following either substance (Fig 4A; Datasets S8 & S9). Inflammation and metabolic-related pathways show a gain of rhythmicity following cocaine or morphine (Figs. 4A & B). For instance, metabolic-related pathways, such as LXR/RXR and FXR/RXR, showed a strong gain in rhythmicity following cocaine. Immune and inflammation-related pathways, such as acute phase response signaling and dendritic cell maturation gain rhythmicity following cocaine or morphine, respectively (Fig. 4A). Moreover, biological processes related to synaptic plasticity (i.e., modulation of synaptic transmission, regulation of long-term synaptic potentiation, dendrite development) showed a gain of rhythmicity following morphine (Fig. 4B).

**Figure 4.**
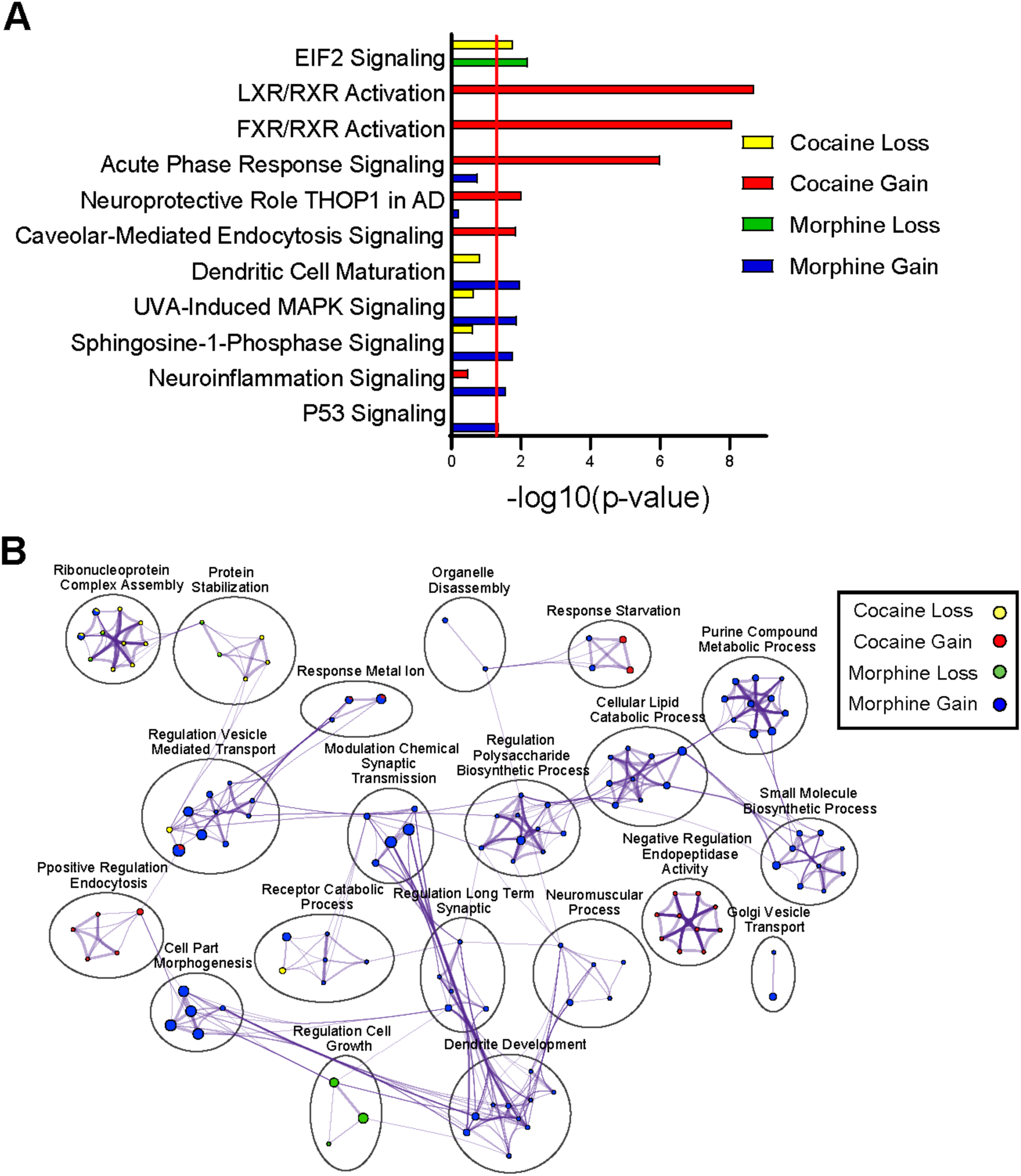
Pathway and biological process enrichment analysis for proteins with a gain or loss of rhythmicity in the NAc following cocaine or morphine. (**A**) Top pathways enriched among the proteins with a gain or loss of rhythmicity in the NAc by drug treatment, as revealed by Ingenuity Pathway Analysis (IPA). A significance cutoff of −Log10(p value) of 1.3, or p<0.05, was used to determine pathway enrichment, depicted by the red line. (**B**) Gene Ontology (GO) Biological process enrichment via Metascape for the top proteins with gained or lost rhythmicity in the NAc by drug treatment. Meta-analysis was used to compare process enrichment across treatment groups by gain or loss of rhythmicity, depicted via a Cytoscape enrichment network plot. Plots were generated using the same parameters as in Figure 2.

### Diurnal rhythms of protein expression following cocaine or morphine contribute to time-of-day-dependent differential expression in NAc

Changes in protein expression following substance use may be associated with altered rhythmicity of the protein. To assess whether diurnal changes in the proteome was related to DE protein, we compared the overlap of DE and rhythmic proteins in NAc within cocaine and morphine groups (Figs. 5A & B; Dataset S10). We found significant overlap between proteins that are rhythmic and DE during the day following cocaine (Fig. 5A). Following morphine, there was overlap between proteins that are rhythmic and DE at night (Fig. 5B). Overall, the overlap between rhythmic and DE proteins following morphine was more robust than cocaine (Fig. 5A, top right panel vs. 5B, bottom right panel). Scatterplots of proteins that overlapped between DE and rhythmicity were associated with both downregulation of protein expression (Fig. 5C) and upregulation (Fig. 5D) particularly during the night, with most proteins displaying downregulation (Dataset S10). Taken together, these data suggest that the rhythmic expression of these proteins following substance administration likely contribute to time-of-day-dependent differential expression of proteins in NAc, suggesting protein alterations following substance use is a dynamic process that depends on diurnal processes.

**Figure 5.**
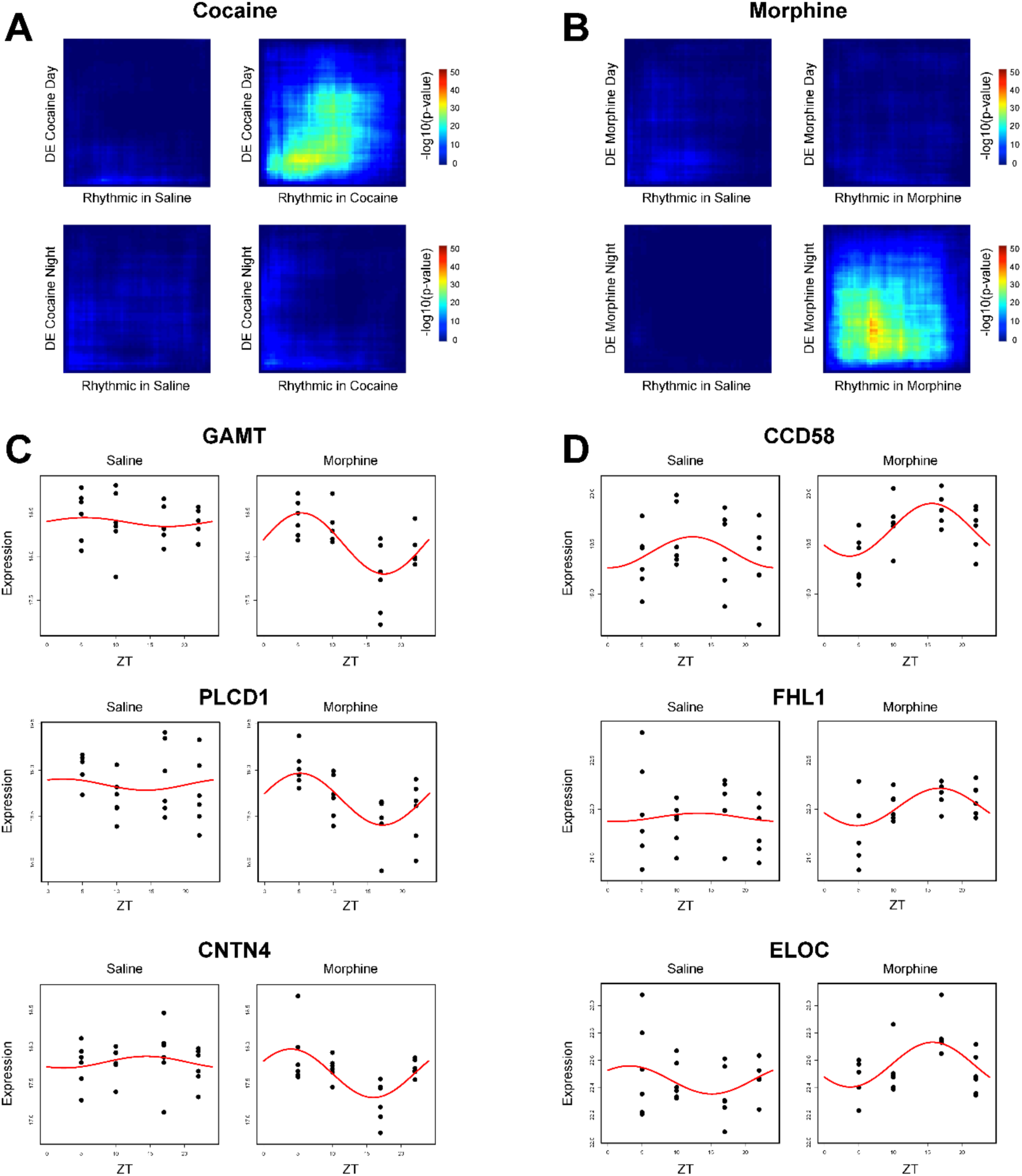
Overlap between rhythmic and DE proteins in the NAc as a result of cocaine or morphine. (**A**) Rank-Rank Hypergeometric Overlap (RRHO) analysis of the overlap between the cocaine-specific rhythmic proteins in the NAc and the DE proteins in the NAc by substance and across time of day. RRHO revealed limited overlap except in the DE Cocaine Day versus Cocaine Rhythmic comparison. (**B**) RRHO analysis of the overlap between the morphine-specific rhythmic proteins in the NAc and the DE proteins in the NAc by substance and across time of day. RRHO revealed limited overlap except in the DE Morphine Night versus Morphine Rhythmic comparison. (**C**) Representative overlapped morphine-specific rhythmic proteins with the largest *downregulation* fold change at night, plotted relative to saline. (**D**) Representative overlapped morphine-specific rhythmic proteins with the largest *upregulation* fold change at night, plotted relative to saline. Dots indicate individual samples, with the y axis indicating expression and x axis indicating time of day (ZT 5, 10, 17, 22). Red line depicts the fitted curve as revealed in the cosiner rhythmicity analysis. Cocaine-specific overlapped rhythmic and DE proteins were not shown due to no proteins meeting significance thresholds.

## DISCUSSION

Substance use significantly alters molecular rhythms in NAc in both humans and rodents ^19,40–45^. To date, studies have largely focused on investigating molecular rhythm changes at the transcript level, such as changes in diurnal expression of core clock genes or even changes in rhythms of expression across the transcriptome. However, given the estimation that only 40% of the variation in a protein’s concentration may be explained by its mRNA abundance ^57,58^, investigating molecular rhythms in the NAc at the *protein*-level may yield greater biological or functional insight. Here, using LC-MS/MS quantitative proteomics, our data reveal significant proteome-wide changes in both phase-dependent DE and the rhythmic regulation of the mouse NAc proteome following cocaine or morphine.

First, looking at DE proteins following drug exposure, cocaine and morphine exerted distinct effects on the NAc proteome that differed across time of day (i.e., day vs. night). This is evidenced by little to no overlap in DE proteins neither between cocaine versus morphine exposure nor between day versus night. Notably, morphine yielded an overall greater number of DE proteins than cocaine, primarily driven by morphine’s augmentation of DE proteins during the night (i.e., nocturnal mouse’s active phase). Interestingly, this is opposite of what has been shown with cocaine’s effects at the transcriptome level. Using bulk RNA-sequencing across six times of day, Brami-Cherrier *et al.* demonstrated cocaine’s effects on rhythmic transcripts in the mouse NAc was largely driven by inducing peaks of expression during the *day* phase (i.e., nocturnal mouse’s inactive phase) ^42^. While a similar study has not been performed with morphine or other opioids in rodents, transcriptional rhythms have been investigated in post-mortem brains of individuals with OUD using the subjects’ times of death as a marker of time of day ^19^. Much like our observation of morphine driving greater DE proteins in the NAc during the mouse’s active phase (i.e., nocturnal night), a greater number of rhythmic transcripts in the NAc of OUD subjects were also found to peak during the active phase (i.e., diurnal day) and *Morphine Addiction* was one of the top enriched biological processes among those transcripts ^19^. However, future investigation is needed into the functional significance of this phase-dependent effect of morphine versus cocaine in the NAc.

Next, when assessing the biological pathways/processes enriched among the drug-induced DE proteins, cocaine and morphine were found to vastly differ in the pathways/processes affected in the NAc. First, regarding cocaine, several stress-response related pathways were enriched among the cocaine-induced DE proteins across both day and night, including *Glucocorticoid Receptor Signaling*, *Corticotropin Releasing Hormone Signaling*, and *Adrenomedullin Signaling*. Though, looking specifically at the night phase, cocaine-induced DE proteins were largely attributed to processes related to *Neurotransmitter Transport* and *Regulation of Axonogenesis*, which exhibited a high degree of interconnectivity in an enrichment network analysis. Together, these data are particularly interesting in that they corroborate the growing body of evidence linking the circadian, stress, and reward systems of the brain ^32,59–64^, as well as the long established relationship between cocaine and changes in neurotransmitter signaling, axonal regulation, and other synaptic plasticity-related processes ^65–67^.

Notably, unlike cocaine, morphine-induced DE proteins were primarily associated with immune and metabolic related pathways/processes across both day and night, including *RXR activation* pathways, *acute phase response signaling*, *CTLA4 signaling in Cytotoxic T Lymphocytes,* and various metabolic-related biological processes. This enrichment of immune related pathways among the morphine-induced DE’s is of particular interest considering the many immune-, inflammation-, and metabolic-related processes enriched among DE transcripts found in postmortem NAc samples of OUD subjects ^4^. Moreover, fentanyl self-administration in Sprague-Dawley rats has also been shown to increase expression of cytokines (e.g., interleukin (IL)1β, IL5), chemokines, tumor necrosis factor α (TNFα), and interferon proteins (e.g., IFNβ and IFNγ) in the NAc ^68^. Our mouse proteomic data add to these previous studies, furthering a need for functional investigation into immune- and metabolic-related mechanisms underlying OUD and potential interactions with the circadian system ^69–72^.

In addition to assessing DE proteins by phase, rhythmicity analyses in the NAc revealed largely distinct rhythmic proteins following cocaine versus morphine exposure, as illustrated by RRHO analysis. Strikingly, morphine induced substantially more gains in rhythmic proteins in the NAc compared to cocaine, with nearly seven times more proteins gaining rhythmicity. Interestingly, although there was nearly no overlap between proteins that gained rhythmicity, there was extensive overlap between proteins that lost rhythmicity following exposure to the two drugs. These observations suggest the two substances may have more in common regarding the way they disrupt rhythms in the NAc, but perhaps differ regarding they ways they reprogram circadian regulation of the NAc – with morphine leading to greater circadian reprogramming than cocaine. While studies in both rodents and human post-mortem tissue have introduced the idea of drugs of abuse inducing circadian reprogramming in the NAc ^19,42^, no studies have directly compared cocaine versus opioids in this context. Future studies should investigate the functional significance of this differential circadian reprogramming by cocaine versus morphine and its role in the development of SUDs. More specifically, expanding to other reward regions of the brain and/or investigating cell-type specific effects of the two drugs may yield greater insight into their differences.

When assessing the biological pathways/processes enriched among the detected proteins that gain or lose rhythmicity, we found proteins related to translation (i.e., EIF2 signaling) similarly lost rhythmicity with cocaine or morphine. This may be related to shared mechanisms by which cocaine or morphine lead to changes in synaptic plasticity to drive their reinforcement ^7,73–75^. Interestingly, synaptic plasticity related pathways/processes were enriched among the top gain of rhythmicity proteins following morphine exposure – something notably not seen among morphine’s DE proteins. This is also true of the RXR activation pathways among the top gain of rhythmicity proteins following cocaine– pathways previously seen among the *morphine*-induced DE proteins. Thus, these observations prompted us to investigate whether rhythmic expression of these proteins following drug exposure could be driving phase-specific DE changes in the NAc.

Strikingly, a phase-dependent differential relationship was found between DE proteins and rhythmic transcripts following cocaine versus morphine. More specifically, rhythmic proteins induced by cocaine exposure primarily overlapped with the cocaine-induced DE proteins during the *day* phase, while rhythmic proteins induced by morphine exposure primarily overlapped with the morphine-induced DE proteins during the *night* phase. Also, morphine showed an overall more, robust overlap when compared to that of cocaine, particularly when significance thresholds were considered. To the best of our knowledge, these data are the first to demonstrate a relationship between drug-induced rhythmic expression of proteins and the induction of phase-specific DE changes in the NAc – with cocaine’s concordance occurring during the day and morphine’s concordance occurring at night. This has major implications for any study assessing DE at single times of day without considering the contribution of circadian / phase-specific effects.

Finally, it is important to note a few limitations in the interpretation of our data. To investigate how cocaine and morphine affect the NAc proteome across time of day, mice were exclusively given either cocaine, morphine, or saline injections. While this allowed us to compare the effects of cocaine versus morphine separately, it does not address how the two substances together may influence the NAc proteome. This would be of particular interest for future studies given the fast growing prevalence of polysubstance use, with opioids commonly used simultaneously with illicit psychostimulants ^76–79^. It is also important to consider our studies only used male mice to study the effects of cocaine versus morphine in NAc. Although the prevalence of substance abuse is significantly greater in males than in females, with twice as many drugs overdose deaths in males than females in 2021 ^80^, females exhibit a more rapid escalation of drug use, greater withdrawal response, and are more vulnerable in terms of treatment outcomes ^81–83^. Moreover, several recent preclinical studies from our lab and others demonstrate significant sex differences in the interaction between the circadian and reward systems that influence reward-related behaviors ^16–18,21,84^. Thus, future studies must investigate the differential circadian effects of cocaine or morphine on the NAc proteome in both males and females ^7,73,85^. Nevertheless, our findings demonstrate the impact of cocaine and morphine on the diurnal rhythmicity of the proteome in the NAc, revealing putatively involved functional consequences on metabolic and neuroinflammatory processes in the brain associated with SUDs.

## ASSOCIATED CONTENT

### Supporting Information

The following Supporting Information is available free of charge:

Dataset S1: List of DE parameters for Fig. 1 (XLSX)

Dataset S2: Filtered list of DE proteins for Figs. 1 and 2 (XLSX)

Dataset S3: Pathway analysis for Fig. 2a (XLSX)

Dataset S4: Metascape analyses for Fig. 2b (XLSX)

Dataset S5: Circadian parameters for Fig. 3 (XLSX)

Dataset S6: Gain/Loss rhythmicity parameters for Fig. 3 (XLSX)

Dataset S7: Filtered list of proteins that gain/loss rhythmicity for Figs. 3 and 4 (XLSX)

Dataset S8: Pathway analysis for Fig. 4a (XLSX)

Dataset S9: Metascape analyses for Fig. 4b (XLSX)

Dataset S10: List of DE and rhythmic overlap for Fig. 5 (XLSX)

## AUTHOR INFORMATION

### Corresponding Author

*. Tel: 508-856-5542

### Author Contributions

‡ KDK and DDBK contributed equally. DBK, RSW, TTL, CAN, KRW, and RWL contributed to the experimental design of this study; DBK and KDK conducted experiments. RSW and TTL performed LC-MS/MS sample preparation, DIA, and DIA analysis under the guidance of KRW and ACN. XX performed biostatistical analyses under the guidance of GCT and RWL. KDK performed pathway enrichment analyses and generated figures. The manuscript was written and edited through contributions of all authors. All authors have given approval to the final version of the manuscript.

### Notes

The authors have no conflict of interest, nor competing financial interest to declare.

## Supporting information

Dataset S1: List of DE parameters for Fig. 1

Dataset S2: Filtered list of DE proteins for Figs. 1 and 2

Dataset S3: Pathway analysis for Fig. 2a

Dataset S4: Metascape analyses for Fig. 2b

Dataset S5: Circadian parameters for Fig. 3

Dataset S6: Gain/Loss rhythmicity parameters for Fig. 3

Dataset S7: Filtered list of proteins that gain/loss rhythmicity for Figs. 3 and 4

Dataset S8: Pathway analysis for Fig. 4a

Dataset S9: Metascape analyses for Fig. 4b

Dataset S10: List of DE and rhythmic overlap for Fig. 5

## ABBREVIATIONS

DDA: Data Dependent Acquisition
DE: Differentially Expressed
DIA: Data independent acquisition
GO: Gene Ontology
LC-MS/MS: liquid chromatography mass spectrometry
IPA: Ingenuity Pathway Analysis
NAc: nucleus accumbens
OUD: Opioid Use Disorder
RNA: ribonucleic acid
RRHO: Rank-Rank Hypergeometric Overlap
SCN: suprachiasmatic nucleus
SUD: Substance Use Disorder
ZT: zeitgeber time

## ACKNOWLEDGMENTS

We thank Weiwei Wang and Florine Collin for their assistance in the LC-MS/MS sample preparation. This work was supported by the National Institutes of Health (NIH) (Yale/NIDA Neuroproteomics Center DA018343 [TTL, KRW, CAN], MH128763 [KDK], DA0386654 [RWL], DA041872 [RWL], and HL150432 [RWL]) and NARSAD Young Investigator Award (Brain & Behavior Foundation; P&S Fund [KDK]). The Orbitrap Fusion mass spectrometer and the offline UPLC was funded in part by NIH SIG from the Office of The Director, National Institutes of Health under Award Numbers (S10OD02365101A1, S10OD018034 and S10OD019967, respectively). The funders had no role in study design, data collection and analysis, decision to publish, or preparation of the manuscript. The content is solely the responsibility of the authors and does not necessarily represent the official views of the National Institutes of Health.

## REFERENCES

(1) Ahmad, F. B.; Rossen, L. M.; Sutton, P. Provisional drug overdose death counts. https://www.cdc.gov/nchs/nvss/vsrr/drug-overdose-data.htm (accessed Feb 8, 2023).

(2) Scofield, M. D.; Heinsbroek, J. A.; Gipson, C. D.; Kupchik, Y. M.; Spencer, S.; Smith, A. C. W.; Roberts-Wolfe, D.; Kalivas, P. W. The Nucleus Accumbens: Mechanisms of Addiction across Drug Classes Reflect the Importance of Glutamate Homeostasis. Pharmacol. Rev. 2016, 68, 816–871.

(3) Walker, D. M.; Cates, H. M.; Loh, Y.-H. E.; Purushothaman, I.; Ramakrishnan, A.; Cahill, K. M.; Lardner, C. K.; Godino, A.; Kronman, H. G.; Rabkin, J.;, et al. Cocaine Self-administration Alters Transcriptome-wide Responses in the Brain’s Reward Circuitry. Biol. Psychiatry 2018, 84, 867–880.

(4) Seney, M. L.; Kim, S.-M.; Glausier, J. R.; Hildebrand, M. A.; Xue, X.; Zong, W.; Wang, J.; Shelton, M. A.; Phan, B. N.; Srinivasan, C.;, et al. Transcriptional alterations in dorsolateral prefrontal cortex and nucleus accumbens implicate neuroinflammation and synaptic remodeling in opioid use disorder. Biol. Psychiatry 2021, 90, 550–562.

(5) Zipperly, M. E.; Sultan, F. A.; Graham, G.-E.; Brane, A. C.; Simpkins, N. A.; Carullo, N. V. N.; Ianov, L.; Day, J. J. Regulation of dopamine-dependent transcription and cocaine action by Gadd45b. Neuropsychopharmacology 2021, 46, 709–720.

(6) Scheyer, A. F.; Wolf, M. E.; Tseng, K. Y. A protein synthesis-dependent mechanism sustains calcium-permeable AMPA receptor transmission in nucleus accumbens synapses during withdrawal from cocaine self-administration. J. Neurosci. 2014, 34, 3095–3100.

(7) Werner, C. T.; Stefanik, M. T.; Milovanovic, M.; Caccamise, A.; Wolf, M. E. Protein Translation in the Nucleus Accumbens Is Dysregulated during Cocaine Withdrawal and Required for Expression of Incubation of Cocaine Craving. J. Neurosci. 2018, 38, 2683–2697.

(8) Luo, J.; Jing, L.; Qin, W.-J.; Zhang, M.; Lawrence, A. J.; Chen, F.; Liang, J.-H. Transcription and protein synthesis inhibitors reduce the induction of behavioural sensitization to a single morphine exposure and regulate Hsp70 expression in the mouse nucleus accumbens. Int. J. Neuropsychopharmacol. 2011, 14, 107–121.

(9) Tannu, N. S.; Howell, L. L.; Hemby, S. E. Integrative proteomic analysis of the nucleus accumbens in rhesus monkeys following cocaine self-administration. Mol. Psychiatry 2010, 15, 185–203.

(10) Zhang, Y.; Crofton, E. J.; Fan, X.; Li, D.; Kong, F.; Sinha, M.; Luxon, B. A.; Spratt, H. M.; Lichti, C. F.; Green, T. A. Convergent transcriptomics and proteomics of environmental enrichment and cocaine identifies novel therapeutic strategies for addiction. Neuroscience 2016, 339, 254–266.

(11) López, A. J.; Johnson, A. R.; Euston, T. J.; Wilson, R.; Nolan, S. O.; Brady, L. J.; Thibeault, K. C.; Kelly, S. J.; Kondev, V.; Melugin, P.;, et al. Cocaine self-administration induces sex-dependent protein expression in the nucleus accumbens. *Commun*. Biol. 2021, 4, 883.

(12) Li, K. W.; Jimenez, C. R.; van der Schors, R. C.; Hornshaw, M. P.; Schoffelmeer, A. N. M.; Smit, A. B. Intermittent administration of morphine alters protein expression in rat nucleus accumbens. Proteomics 2006, 6, 2003–2008.

(13) Van den Oever, M. C.; Lubbers, B. R.; Goriounova, N. A.; Li, K. W.; Van der Schors, R. C.; Loos, M.; Riga, D.; Wiskerke, J.; Binnekade, R.; Stegeman, M.;, et al. Extracellular matrix plasticity and GABAergic inhibition of prefrontal cortex pyramidal cells facilitates relapse to heroin seeking. Neuropsychopharmacology 2010, 35, 2120–2133.

(14) Bu, Q.; Yang, Y.; Yan, G.; Hu, Z.; Hu, C.; Duan, J.; Lv, L.; Zhou, J.; Zhao, J.; Shao, X.;, et al. Proteomic analysis of the nucleus accumbens in rhesus monkeys of morphine dependence and withdrawal intervention. J. Proteomics 2012, 75, 1330–1342.

(15) Park, H.-M.; Satta, R.; Davis, R. G.; Goo, Y. A.; LeDuc, R. D.; Fellers, R. T.; Greer, J. B.; Romanova, E. V.; Rubakhin, S. S.; Tai, R.;, et al. Multidimensional Top-Down Proteomics of Brain-Region-Specific Mouse Brain Proteoforms Responsive to Cocaine and Estradiol. J. Proteome Res. 2019, 18, 3999–4012.

(16) Puig, S.; Shelton, M. A.; Barko, K.; Seney, M. L.; Logan, R. W. Sex-specific role of the circadian transcription factor NPAS2 in opioid tolerance, withdrawal and analgesia. Genes Brain Behav. 2022, 21, e12829.

(17) Gamble, M. C.; Chuan, B.; Gallego-Martin, T.; Shelton, M. A.; Puig, S.; O’Donnell, C. P.; Logan, R. W. A role for the circadian transcription factor NPAS2 in the progressive loss of non-rapid eye movement sleep and increased arousal during fentanyl withdrawal in male mice. Psychopharmacology 2022.

(18) Becker-Krail, D. D.; Ketchesin, K. D.; Burns, J. N.; Zong, W.; Hildebrand, M. A.; DePoy, L. M.; Vadnie, C. A.; Tseng, G. C.; Logan, R. W.; Huang, Y. H.;, et al. Astrocyte Molecular Clock Function in the Nucleus Accumbens is Important for Reward-Related Behavior. Biol. Psychiatry 2022.

(19) Xue, X.; Zong, W.; Glausier, J. R.; Kim, S.-M.; Shelton, M. A.; Phan, B. N.; Srinivasan, C.; Pfenning, A. R.; Tseng, G. C.; Lewis, D. A.;, et al. Molecular rhythm alterations in prefrontal cortex and nucleus accumbens associated with opioid use disorder. Transl. Psychiatry 2022, 12, 123.

(20) Becker-Krail, D. D.; Parekh, P. K.; Ketchesin, K. D.; Yamaguchi, S.; Yoshino, J.; Hildebrand, M. A.; Dunham, B.; Ganapathiraju, M. K.; Logan, R. W.; McClung, C. A. Circadian transcription factor NPAS2 and the NAD+-dependent deacetylase SIRT1 interact in the mouse nucleus accumbens and regulate reward. Eur. J. Neurosci. 2022, 55, 675–693.

(21) DePoy, L. M.; Becker-Krail, D. D.; Zong, W.; Petersen, K.; Shah, N. M.; Brandon, J. H.; Miguelino, A. M.; Tseng, G. C.; Logan, R. W.; McClung, C. A. Circadian-Dependent and Sex-Dependent Increases in Intravenous Cocaine Self-Administration in Npas2 Mutant Mice. J. Neurosci. 2021, 41, 1046–1058.

(22) Logan, R. W.; McClung, C. A. Rhythms of life: circadian disruption and brain disorders across the lifespan. Nat. Rev. Neurosci. 2019, 20, 49–65.

(23) Logan, R. W.; Parekh, P. K.; Kaplan, G. N.; Becker-Krail, D. D.; Williams, W. P.; Yamaguchi, S.; Yoshino, J.; Shelton, M. A.; Zhu, X.; Zhang, H.;, et al. NAD+ cellular redox and SIRT1 regulate the diurnal rhythms of tyrosine hydroxylase and conditioned cocaine reward. Mol. Psychiatry 2019, 24, 1668–1684.

(24) Logan, R. W.; Hasler, B. P.; Forbes, E. E.; Franzen, P. L.; Torregrossa, M. M.; Huang, Y. H.; Buysse, D. J.; Clark, D. B.; McClung, C. A. Impact of sleep and circadian rhythms on addiction vulnerability in adolescents. Biol. Psychiatry 2018, 83, 987–996.

(25) Eacret, D.; Veasey, S. C.; Blendy, J. A. Bidirectional Relationship between Opioids and Disrupted Sleep: Putative Mechanisms. Mol. Pharmacol. 2020, 98, 445–453.

(26) Eacret, D.; Lemchi, C.; Caulfield, J. I.; Cavigelli, S. A.; Veasey, S. C.; Blendy, J. A. Chronic sleep deprivation blocks voluntary morphine consumption but not conditioned place preference in mice. Front. Neurosci. 2022, 16, 836693.

(27) Hastings, M. H.; Maywood, E. S.; Brancaccio, M. Generation of circadian rhythms in the suprachiasmatic nucleus. Nat. Rev. Neurosci. 2018, 19, 453–469.

(28) Mohawk, J. A.; Green, C. B.; Takahashi, J. S. Central and peripheral circadian clocks in mammals. Annu. Rev. Neurosci. 2012, 35, 445–462.

(29) Begemann, K.; Neumann, A.-M.; Oster, H. Regulation and function of extra-SCN circadian oscillators in the brain. Acta Physiol (Oxf) 2020, 229, e13446.

(30) Becker-Krail, D. D.; Walker, W. H.; Nelson, R. J. The ventral tegmental area and nucleus accumbens as circadian oscillators: implications for drug abuse and substance use disorders. Front. Physiol. 2022, 13, 886704.

(31) Partch, C. L.; Green, C. B.; Takahashi, J. S. Molecular architecture of the mammalian circadian clock. Trends Cell Biol. 2014, 24, 90–99.

(32) Logan, R. W.; Edgar, N.; Gillman, A. G.; Hoffman, D.; Zhu, X.; McClung, C. A. Chronic Stress Induces Brain Region-Specific Alterations of Molecular Rhythms that Correlate with Depression-like Behavior in Mice. Biol. Psychiatry 2015, 78, 249–258.

(33) Ozburn, A. R.; Kern, J.; Parekh, P. K.; Logan, R. W.; Liu, Z.; Falcon, E.; Becker-Krail, D.; Purohit, K.; Edgar, N. M.; Huang, Y.;, et al. NPAS2 Regulation of Anxiety-Like Behavior and GABAA Receptors. Front. Mol. Neurosci. 2017, 10, 360.

(34) Spencer, S.; Falcon, E.; Kumar, J.; Krishnan, V.; Mukherjee, S.; Birnbaum, S. G.; McClung, C. A. Circadian genes Period 1 and Period 2 in the nucleus accumbens regulate anxiety-related behavior. Eur. J. Neurosci. 2013, 37, 242–250.

(35) Zhao, C.; Gammie, S. C. The circadian gene Nr1d1 in the mouse nucleus accumbens modulates sociability and anxiety-related behaviour. Eur. J. Neurosci. 2018, 48, 1924–1943.

(36) Parekh, P. K.; Logan, R. W.; Ketchesin, K. D.; Becker-Krail, D.; Shelton, M. A.; Hildebrand, M. A.; Barko, K.; Huang, Y. H.; McClung, C. A. Cell-Type-Specific Regulation of Nucleus Accumbens Synaptic Plasticity and Cocaine Reward Sensitivity by the Circadian Protein, NPAS2. J. Neurosci. 2019, 39, 4657–4667.

(37) Ozburn, A. R.; Falcon, E.; Twaddle, A.; Nugent, A. L.; Gillman, A. G.; Spencer, S. M.; Arey, R. N.; Mukherjee, S.; Lyons-Weiler, J.; Self, D. W.;, et al. Direct regulation of diurnal Drd3 expression and cocaine reward by NPAS2. Biol. Psychiatry 2015, 77, 425–433.

(38) Porcu, A.; Vaughan, M.; Nilsson, A.; Arimoto, N.; Lamia, K.; Welsh, D. K. Vulnerability to helpless behavior is regulated by the circadian clock component CRYPTOCHROME in the mouse nucleus accumbens. Proc. Natl. Acad. Sci. USA 2020, 117, 13771–13782.

(39) Falcon, E.; Ozburn, A.; Mukherjee, S.; Roybal, K.; McClung, C. A. Differential regulation of the period genes in striatal regions following cocaine exposure. PLoS One 2013, 8, e66438.

(40) Wang, D.-Q.; Wang, X.-L.; Wang, C.-Y.; Wang, Y.; Li, S.-X.; Liu, K.-Z. Effects of chronic cocaine exposure on the circadian rhythmic expression of the clock genes in reward-related brain areas in rats. Behav. Brain Res. 2019, 363, 61–69.

(41) Brami-Cherrier, K.; Lewis, R. G.; Cervantes, M.; Liu, Y.; Tognini, P.; Baldi, P.; Sassone-Corsi, P.; Borrelli, E. Cocaine-mediated circadian reprogramming in the striatum through dopamine D2R and PPARγ activation. Nat. Commun. 2020, 11, 4448.

(42) Li, S.; Liu, L.; Jiang, W.; Lu, L. Morphine withdrawal produces circadian rhythm alterations of clock genes in mesolimbic brain areas and peripheral blood mononuclear cells in rats. J. Neurochem. 2009, 109, 1668–1679.

(43) Li, S.; Liu, L.; Jiang, W.; Sun, L.; Zhou, S.; Le Foll, B.; Zhang, X. Y.; Kosten, T. R.; Lu, L. Circadian alteration in neurobiology during protracted opiate withdrawal in rats. J. Neurochem. 2010, 115, 353–362.

(44) Zhang, P.; Moye, L. S.; Southey, B. R.; Dripps, I.; Sweedler, J. V.; Pradhan, A.; Rodriguez-Zas, S. L. Opioid-Induced Hyperalgesia Is Associated with Dysregulation of Circadian Rhythm and Adaptive Immune Pathways in the Mouse Trigeminal Ganglia and Nucleus Accumbens. Mol. Neurobiol. 2019, 56, 7929–7949.

(45) Castañeda, T. R.; de Prado, B. M.; Prieto, D.; Mora, F. Circadian rhythms of dopamine, glutamate and GABA in the striatum and nucleus accumbens of the awake rat: modulation by light. J Pineal Res 2004, 36, 177–185.

(46) Baltazar, R. M.; Coolen, L. M.; Webb, I. C. Diurnal rhythms in neural activation in the mesolimbic reward system: critical role of the medial prefrontal cortex. Eur. J. Neurosci. 2013, 38, 2319–2327.

(47) Ketchesin, K. D.; Zong, W.; Hildebrand, M. A.; Seney, M. L.; Cahill, K. M.; Scott, M. R.; Shankar, V. G.; Glausier, J. R.; Lewis, D. A.; Tseng, G. C.;, et al. Diurnal rhythms across the human dorsal and ventral striatum. Proc. Natl. Acad. Sci. USA 2021, 118.

(48) Stahl, D. C.; Swiderek, K. M.; Davis, M. T.; Lee, T. D. Data-controlled automation of liquid chromatography/tandem mass spectrometry analysis of peptide mixtures. J Am Soc Mass Spectrom 1996, 7, 532–540.

(49) Venable, J. D.; Dong, M.-Q.; Wohlschlegel, J.; Dillin, A.; Yates, J. R. Automated approach for quantitative analysis of complex peptide mixtures from tandem mass spectra. Nat. Methods 2004, 1, 39–45.

(50) Gillet, L. C.; Navarro, P.; Tate, S.; Röst, H.; Selevsek, N.; Reiter, L.; Bonner, R.; Aebersold, R. Targeted data extraction of the MS/MS spectra generated by data-independent acquisition: a new concept for consistent and accurate proteome analysis. Mol. Cell Proteomics 2012, 11, O111.016717.

(51) Collins, B. C.; Hunter, C. L.; Liu, Y.; Schilling, B.; Rosenberger, G.; Bader, S. L.; Chan, D. W.; Gibson, B. W.; Gingras, A.-C.; Held, J. M.;, et al. Multi-laboratory assessment of reproducibility, qualitative and quantitative performance of SWATH-mass spectrometry. Nat. Commun. 2017, 8, 291.

(52) Searle, B. C.; Pino, L. K.; Egertson, J. D.; Ting, Y. S.; Lawrence, R. T.; MacLean, B. X.; Villén, J.; MacCoss, M. J. Chromatogram libraries improve peptide detection and quantification by data independent acquisition mass spectrometry. Nat. Commun. 2018, 9, 5128.

(53) Krämer, A.; Green, J.; Pollard, J.; Tugendreich, S. Causal analysis approaches in Ingenuity Pathway Analysis. Bioinformatics 2014, 30, 523–530.

(54) Zhou, Y.; Zhou, B.; Pache, L.; Chang, M.; Khodabakhshi, A. H.; Tanaseichuk, O.; Benner, C.; Chanda, S. K. Metascape provides a biologist-oriented resource for the analysis of systems-level datasets. Nat. Commun. 2019, 10, 1523.

(55) Morrison, K. E.; Stenson, A. F.; Marx-Rattner, R.; Carter, S.; Michopoulos, V.; Gillespie, C. F.; Powers, A.; Huang, W.; Kane, M. A.; Jovanovic, T.;, et al. Developmental timing of trauma in women predicts unique extracellular vesicle proteome signatures. Biol. Psychiatry 2022, 91, 273–282.

(56) Vogel, C.; Marcotte, E. M. Insights into the regulation of protein abundance from proteomic and transcriptomic analyses. Nat. Rev. Genet. 2012, 13, 227–232.

(57) Liu, Y.; Beyer, A.; Aebersold, R. On the Dependency of Cellular Protein Levels on mRNA Abundance. Cell 2016, 165, 535–550.

(58) Becker-Krail, D.; McClung, C. Implications of circadian rhythm and stress in addiction vulnerability. [version 1; peer review: 2 approved]. F1000Res. 2016, 5, 59.

(59) Logan, R. W.; Williams, W. P.; McClung, C. A. Circadian rhythms and addiction: mechanistic insights and future directions. Behav. Neurosci. 2014, 128, 387–412.

(60) Koch, C. E.; Leinweber, B.; Drengberg, B. C.; Blaum, C.; Oster, H. Interaction between circadian rhythms and stress. Neurobiol. Stress 2017, 6, 57–67.

(61) Landgraf, D.; McCarthy, M. J.; Welsh, D. K. Circadian clock and stress interactions in the molecular biology of psychiatric disorders. Curr Psychiatry Rep 2014, 16, 483.

(62) Perreau-Lenz, S.; Spanagel, R. Clock genes × stress × reward interactions in alcohol and substance use disorders. Alcohol 2015, 49, 351–357.

(63) Baik, J.-H. Stress and the dopaminergic reward system. Exp Mol Med 2020, 52, 1879–1890.

(64) Wolf, M. E. Synaptic mechanisms underlying persistent cocaine craving. Nat. Rev. Neurosci. 2016, 17, 351–365.

(65) Lüscher, C.; Malenka, R. C. Drug-evoked synaptic plasticity in addiction: from molecular changes to circuit remodeling. Neuron 2011, 69, 650–663.

(66) Creed, M. C.; Lüscher, C. Drug-evoked synaptic plasticity: beyond metaplasticity. Curr. Opin. Neurobiol. 2013, 23, 553–558.

(67) Ezeomah, C.; Cunningham, K. A.; Stutz, S. J.; Fox, R. G.; Bukreyeva, N.; Dineley, K. T.; Paessler, S.; Cisneros, I. E. Fentanyl self-administration impacts brain immune responses in male Sprague-Dawley rats. Brain Behav. Immun. 2020, 87, 725–738.

(68) Neuroinflammation, Metabolism, and Psychiatric Disorders; Freyberg, Z.; Logan, R. W.; Leboyer, M.; Penninx, B., Eds.; Frontiers Media SA, 2022.

(69) Logan, R. W.; Sarkar, D. K. Circadian nature of immune function. Mol. Cell. Endocrinol. 2012, 349, 82–90.

(70) Labrecque, N.; Cermakian, N. Circadian clocks in the immune system. J. Biol. Rhythms 2015, 30, 277–290.

(71) Freyberg, Z.; Logan, R. W. The intertwined roles of circadian rhythms and neuronal metabolism fueling drug reward and addiction. Curr. Opin. Physiol. 2018, 5, 80–89.

(72) Jian, M.; Luo, Y.-X.; Xue, Y.-X.; Han, Y.; Shi, H.-S.; Liu, J.-F.; Yan, W.; Wu, P.; Meng, S.-Q.; Deng, J.-H.;, et al. eIF2α dephosphorylation in basolateral amygdala mediates reconsolidation of drug memory. J. Neurosci. 2014, 34, 10010–10021.

(73) Placzek, A. N.; Prisco, G. V. D.; Khatiwada, S.; Sgritta, M.; Huang, W.; Krnjević, K.; Kaufman, R. J.; Dani, J. A.; Walter, P.; Costa-Mattioli, M. eIF2α-mediated translational control regulates the persistence of cocaine-induced LTP in midbrain dopamine neurons. Elife 2016, 5.

(74) Laguesse, S.; Ron, D. Protein translation and psychiatric disorders. Neuroscientist 2020, 26, 21–42.

(75) Cicero, T. J.; Ellis, M. S.; Kasper, Z. A. Polysubstance use: A broader understanding of substance use during the opioid crisis. Am. J. Public Health 2020, 110, 244–250.

(76) Ellis, M. S.; Kasper, Z. A.; Cicero, T. J. Polysubstance use trends and variability among individuals with opioid use disorder in rural versus urban settings. Prev. Med. 2021, 152, 106729.

(77) Strickland, J. C.; Havens, J. R.; Stoops, W. W. A nationally representative analysis of “twin epidemics”: Rising rates of methamphetamine use among persons who use opioids. Drug Alcohol Depend. 2019, 204, 107592.

(78) Shearer, R. D.; Jones, A.; Howell, B. A.; Segel, J. E.; Winkelman, T. N. A. Associations between prescription and illicit stimulant and opioid use in the United States, 2015-2020. J Subst Abuse Treat 2022, 143, 108894.

(79) Spencer, M.; Miniño, A.; Warner, M. Drug overdose deaths in the united states, 2001–2021; Centers for Disease Control and Prevention: Atlanta, Georgia, 2022.

(80) NIDA. Drug Overdose Death Rates | National Institute on Drug Abuse (NIDA) https://nida.nih.gov/research-topics/trends-statistics/overdose-death-rates (accessed Feb 14, 2023).

(81) Becker, J. B. Sex differences in addiction. Dialogues Clin Neurosci 2016, 18, 395–402.

(82) Becker, J. B.; Chartoff, E. Sex differences in neural mechanisms mediating reward and addiction. Neuropsychopharmacology 2019, 44, 166–183.

(83) Puig, S.; Shelton, M. A.; Logan, R. W. The Circadian Transcription Factor NPAS2 Modulates Fentanyl-mediated Withdrawal and Hyperalgesia but Not Tolerance, in a Sex-dependent Manner. J. Pain 2022, 23, 59.

(84) Alcantara, A. A.; Lim, H. Y.; Floyd, C. E.; Garces, J.; Mendenhall, J. M.; Lyons, C. L.; Berlanga, M. L. Cocaine- and morphine-induced synaptic plasticity in the nucleus accumbens. Synapse 2011, 65, 309–320.

